# A phenome-wide approach to identify causal risk factors for deep vein thrombosis

**DOI:** 10.1101/476135

**Authors:** Andrei-Emil Constantinescu, Caroline J Bull, Lucy J Goudswaard, Jie Zheng, Benjamin Elsworth, Nicholas J Timpson, Samantha F Moore, Ingeborg Hers, Emma E Vincent

**Affiliations:** MRC Integrative Epidemiology Unit at the University of Bristol, Bristol, UK; Bristol Medical School, Population Health Sciences, University of Bristol, Bristol, UK; School of Translational Health Sciences, Bristol Medical School, University of Bristol, Bristol, United Kingdom; School of Physiology, Pharmacology and Neuroscience, University of Bristol, Bristol, United Kingdom; Department of Endocrine and Metabolic Diseases, Shanghai Institute of Endocrine and Metabolic Diseases, Ruijin Hospital, Shanghai Jiao Tong University School of Medicine, Shanghai, China; Shanghai National Clinical Research Center for Metabolic Diseases, Key Laboratory for Endocrine and Metabolic Diseases of the National Health Commission of the PR China, Shanghai National Center for Translational Medicine, Ruijin Hospital, Shanghai Jiao Tong University School of Medicine, Shanghai, China; Our Future Health. Registered office: 2 New Bailey, 6 Stanley Street, Manchester M3 5GS.

**Keywords:** Mendelian randomization, Deep vein thrombosis, ALSPAC, pQTL, GWAS

## Abstract

**Background:** Deep vein thrombosis (DVT) is the formation of a blood clot in a deep vein. DVT can lead to a venous thromboembolism (VTE), the combined term for DVT and pulmonary embolism, a leading cause of death and disability worldwide. Despite the prevalence and associated morbidity of DVT, the underlying causes are not well understood.

**Objective:** To leverage publicly available genetic summary association statistics to identify causal risk factors for DVT.

**Methods & Results:** We conducted a Mendelian randomization phenome-wide association study (MR-PheWAS) using genetic summary association statistics for 973 exposures and DVT (6,767 cases and 330,392 controls in UK Biobank). There was evidence for a causal effect of 57 exposures on DVT risk, including previously reported risk factors (e.g. body mass index - BMI and height) and novel risk factors (e.g. hyperthyroidism, chronic obstructive pulmonary disease (COPD) and varicose veins). As the majority of identified risk factors were adiposity-related, we explored the molecular link with DVT by undertaking a two-sample MR mediation analysis of BMI-associated circulating proteins on DVT risk. Our results indicate that circulating neurogenic locus notch homolog protein 1 (NOTCH1), inhibin beta C chain (INHBC) and plasminogen activator inhibitor 1 (PAI-1) influence DVT risk, with PAI-1 mediating the BMI-DVT relationship.

**Conclusion:** Using a phenome-wide approach, we provide putative causal evidence that hyperthyroidism, varicose veins, COPD and BMI enhance the risk of DVT. The circulating protein PAI-1 has furthermore a causal role in DVT aetiology and is involved in mediating the BMI-DVT relationship.

## Introduction

Under normal physiological conditions, platelets and fibrin form clots to prevent blood loss at the site of vessel injury [1]. However, when clots (or thromboses) form abnormally they can disrupt blood flow [2,3] and when this occurs in the deep veins of the limbs or pelvis this is known as deep vein thrombosis (DVT). A complication of DVT is pulmonary embolism (PE), where a clot breaks away from a deep vein wall and becomes lodged in a pulmonary blood vessel, obstructing blood flow to the lungs and causing respiratory dysfunction. In 2021, there were approximately one million incident cases of venous thromboembolism (VTE) in the United states alone [4]. DVT accounts for approximately two-thirds of VTE events and PE is the primary contributor to mortality. While VTE was a primary cause for 10,511 deaths in the UK in 2020 [5], the actual contribution of VTE to annual deaths is estimated to be 2-3 fold higher [6].

To prevent acute and chronic complications it is essential to establish an accurate diagnosis of DVT. The symptoms of DVT alone are often not specific or sufficient to make a diagnosis, and about half of those suffering DVT will have no symptoms [7]. Symptoms are considered in conjunction with known risk factors to help estimate the likelihood of DVT and determine whether thromboprophylaxis is required [3]. Pharmacological thromboprophylaxis includes the use of anticoagulants, such as intravenous heparin and oral warfarin (a vitamin K antagonist), which have been used in combination to treat DVT for over 50 years, but require constant maintenance and monitoring [3]. More recently direct oral anticoagulants (DOAC), such as dabigatran (which inhibits thrombin) or rivaroxaban (which inhibits factor Xa), have been employed with reduced economic costs relative to traditional treatments [8].

Risk factors for DVT include genetic factors, such as deficiencies in the anticoagulation proteins antithrombin, protein C, and protein S, or acquired factors, such as age, obesity and smoking [2,9,10]. However, the mechanisms through which these risk factors act have not been clearly established. The identification of novel causal risk factors and potential drug targets is required for improved DVT prophylaxis [3].

Mendelian randomization (MR) allows us to infer causality while addressing limitations of observational epidemiology such as confounding and reverse causation [11–14]. The design of a MR analysis is analogous to that of a randomised control trial (RCT), the “gold standard” method for evaluating the effectiveness of an intervention (**Supplementary Figure 1**) [15]. It is an instrumental variable-based method that uses genetic variants as proxies (or instruments) for exposures to permit causal inference when interpreting relationships between these exposures and disease outcomes [16]. Here, we have used 2-sample MR, which uses data from separate genome-wide association studies (GWAS) for exposures and outcomes of interest [17] to consider the effect of multiple exposures (phenotypes) on DVT risk.

Our MR phenome-wide association study (MR-PheWAS) identified novel risk factors for DVT (e.g. hyperthyroidism and varicose veins) and provided evidence of causality for several previously identified traits (e.g. BMI and height). Of 57 exposures yielding estimates of causal effect on DVT risk, 24 were adiposity-related. While adiposity is an established risk factor for DVT, the biological mechanisms underlying the effect of adiposity and DVT are not well understood. To investigate this mechanistic relationship, we explored whether levels of circulating proteins, known to be altered by adiposity, were in part responsible for this association. A two-sample MR mediation analysis revealed plasminogen activator inhibitor 1 (PAI-1) as a mediator for this relationship.

## Methods

### Study design

With the aim to identify novel risk factors for DVT, we performed an MR-PheWAS to estimate the effects of 973 exposures on DVT risk. As 24 of the 57 exposures that we found causal evidence for an association with DVT were adiposity-related traits (see **Table 1)**, we next decided to investigate potential mediators of this mechanistic relationship further. As previous MR studies have found that levels of circulating proteins are altered by adiposity [18,19], we performed a two-sample mediation MR to estimate the effect of BMI on DVT with BMI-associated proteins as mediators. An overview of the study design is shown in **Figure 1**.

**Figure 1.**
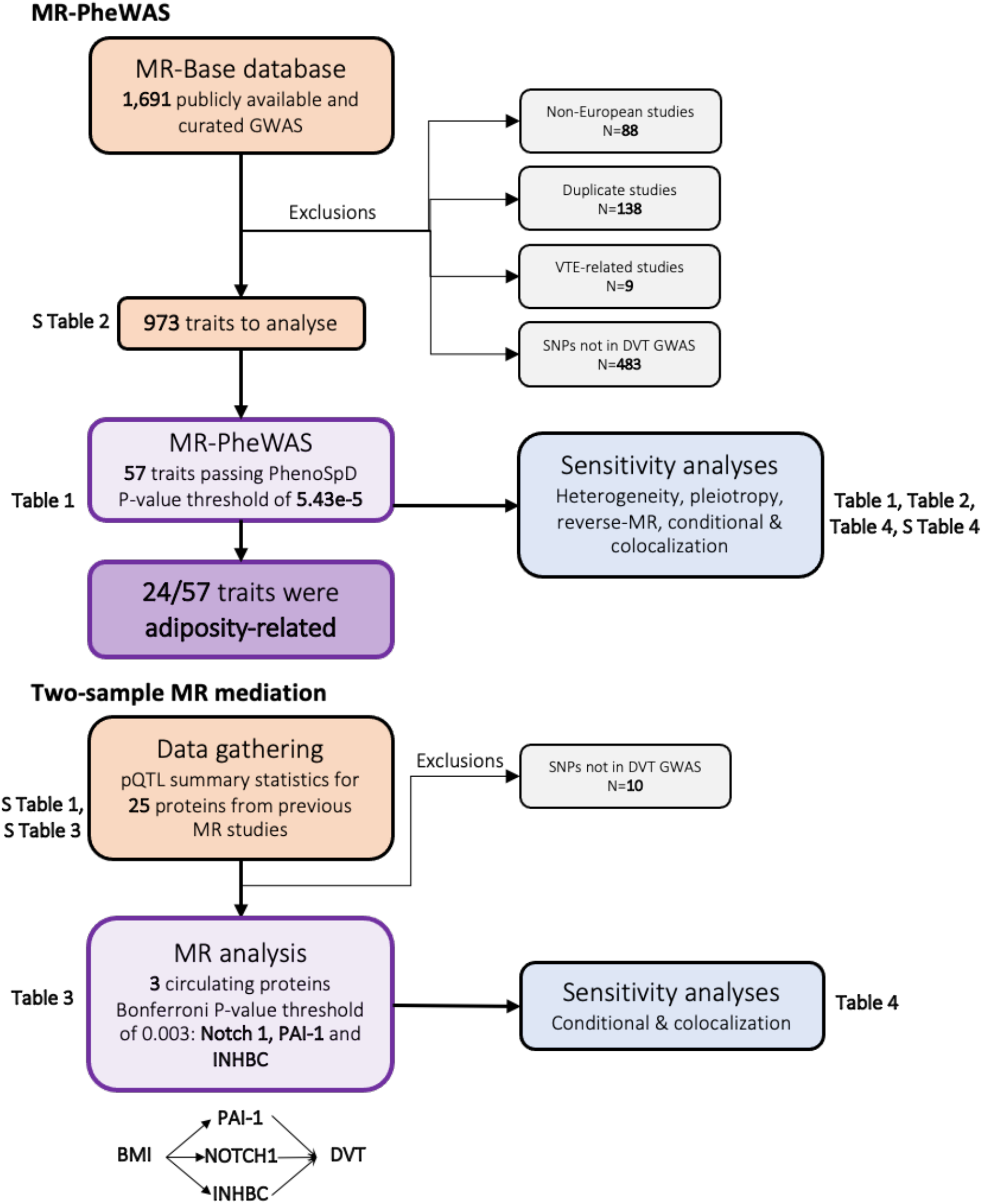
Overview of the study. First, a MR-PheWAS analysis to find risk factors for DVT was done using the MR-Base database and identified many of these to be associated with adiposity (N=24/57). This was followed by a two-sample mediation MR between BMI-associated pQTL data on DVT risk. MR = mendelian randomization; GWAS = genome-wide association study; VTE = venous thromboembolism; DVT = deep vein thrombosis; SNP = single-nucleotide polymorphism; pQTL = protein quantitative trait loci; PAI-1 = Plasminogen activator inhibitor-1; NOTCH1 = Neurogenic locus notch homolog protein 1; INHBC = Inhibin Subunit Beta C; S Table = Supplementary Table.

**Table 1.**
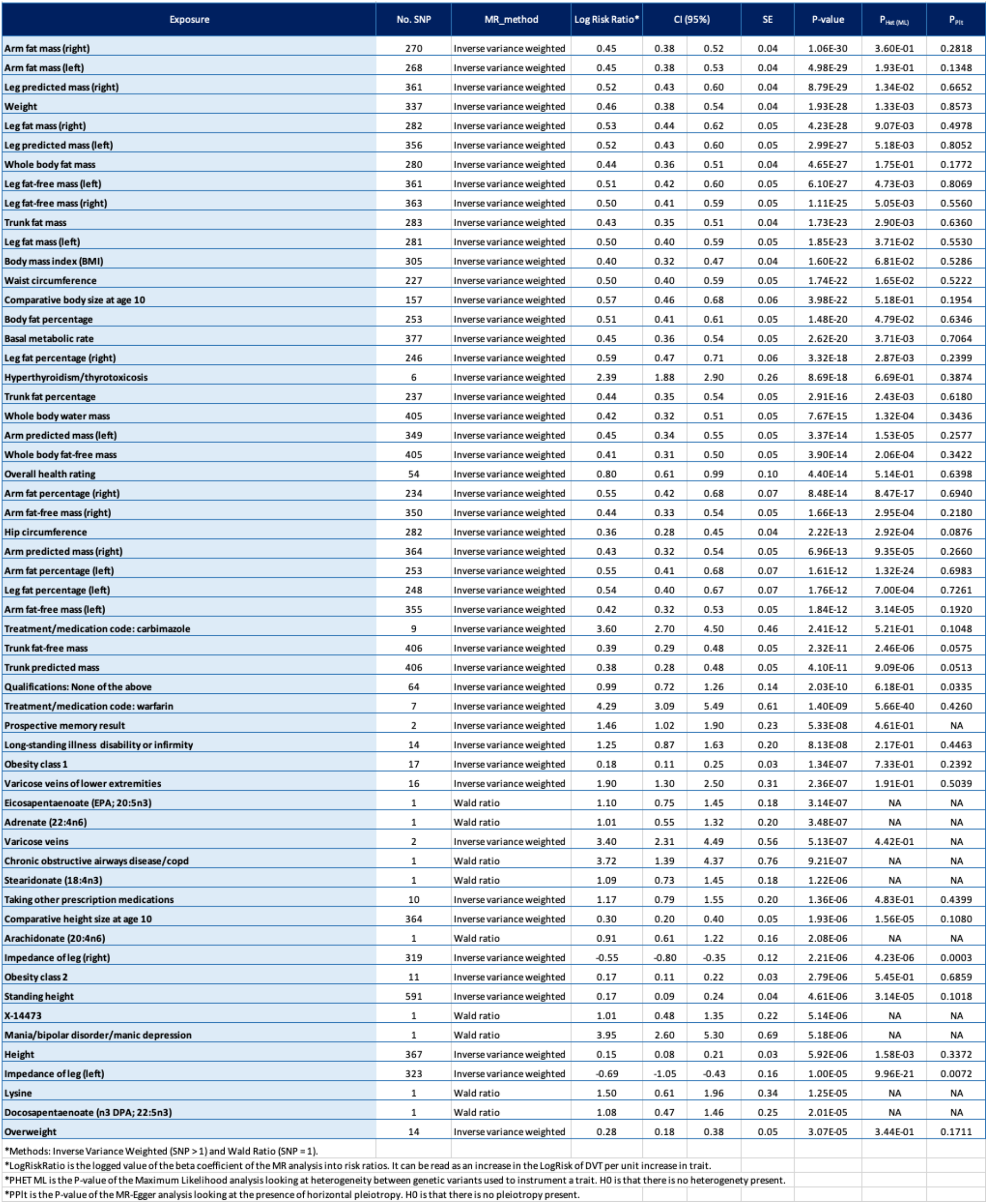
Traits passing the PhenoSpD significance threshold (5.43E-5) in the MR-PheWAS of all traits in UK Biobank on DVT risk.

### Data preparation

#### GWAS data for exposures

All analyses were conducted using R version 3.6.1. The MR-PheWAS was conducted using the TwoSampleMR R package [14]. Genetic data for exposures were obtained from the MR-Base platform of harmonised GWAS summary data [14]. The MR-Base platform permits the hypothesis-free analysis of all catalogued exposures to DVT. The exposures encompassed lifestyle (e.g. BMI and education), disease (e.g. ulcerative colitis and squamous cell carcinoma) and biological (e.g. bone density and oestrogen levels) traits from resources such as UK Biobank [20]. A list of studies with available GWAS summary statistics was obtained through the MR-Base API in R Studio. Non-European (N=88) and duplicate (N=138) studies were excluded. In the case of duplicate studies, those with the highest sample size were retained. VTE (DVT and PE) and VTE-related (e.g. phlebitis and thrombophlebitis) traits were removed (N=9). The genetic instruments used for the analysis were singlenucleotide polymorphisms (SNPs) associated with each of the exposures at a genome-wide level of significance (P<5e-8). As genetic confounding may bias MR estimates if SNPs are correlated [21], linkage disequilibrium (LD) clumping in PLINK [22] was conducted to ensure the SNPs used to instrument exposures were independent (radius = 10,000kb; r^2^ = 0.001) using the 1000 Genomes European reference panel [23]. We also used the 1000 Genomes European dataset [23] to identify potential SNP proxies (with which the initial SNP is in LD with, r^2^>0.8) for those SNPs not present in the DVT data. Depending on the nature of the exposure, the reported effect size for a given SNP was expressed along with the standard error (SE) as a one standard deviation (SD) increase in the level of the risk factor for a continuous exposure, or as a unit change in the exposure on the log-odds scale for a binary trait. Although the number of traits in MR-Base is large and continues to grow, we were limited by the traits available at the time of the analysis. Moreover, some of our exposures did not have a SNP or proxy present in the outcome (DVT) dataset, making it not feasible to perform MR analysis.

#### Deep vein thrombosis GWAS data

Our outcome of interest (DVT) was presented in MR-Base as “Non-cancer illness code self-reported: deep venous thrombosis (dvt)”; these summary results describe a GWAS of 6,767 cases and 330,392 controls done in Europeans. These data describe results of the Neale Lab analysis of UK Biobank data [20] using the PHEnome Scan ANalysis Tool (PHESANT), followed by genotypic data selected through SNP quality control (QC) [24] (https://github.com/Nealelab/UK_Biobank_GWAS).

#### Protein quantitative trait locus data

We aimed to determine whether BMI-associated proteins were mediating the relationship between adiposity and DVT. Our list of BMI-associated proteins was obtained from two previous MR studies investigating the effect of BMI on the circulating proteome [18,19]. We used pQTL data to identify SNPs associated with circulating protein levels at a genome wide level of significance (P ≤ 5e-08). Protein detection platforms for the pQTL data included the SOMAScan^®^ by SomaLogic and Olink (ProSeek CVD array I) [25–28]. Twenty-five proteins were identified using these criteria (**Supplementary Table 1**). PLINK clumping (radius = 10,000kb; r^2^ = 0.001) was performed to ensure the genetic variants used to instrument protein levels were independent. Proxy SNPs for those SNPs that were not present in the DVT data were identified through the 1000 Genomes European dataset [23].

#### Data harmonisation

The majority of GWAS present the effects of a SNP on a trait in relation to the allele on the forward strand. However, the allele present on the forward strand can change over time as reference panels get updated. This requires correction (harmonisation) so that both exposure and outcome data reference the same strand [29]. For exposure and outcome data harmonisation, incorrect but unambiguous alleles were corrected, while ambiguous alleles were removed. In the case of palindromic SNPs (A/T or C/G), allele frequencies were used to solve ambiguities. Harmonisation was not possible for 483 exposures (variants were not present in the DVT GWAS), resulting in a final list of 973 exposures to include in the MR-PheWAS (**Supplementary Table 2**). For our pQTL analysis, 15 of 25 proteins had genetic variants (including proxies) available in the DVT GWAS (**Supplementary Table 3**).

### Mendelian Randomization Analyses

#### MR-PheWAS

A hypothesis-free MR-PheWAS was conducted using the TwoSampleMR R package [30]. The effect of a given exposure on DVT was estimated using the inverse-variance weighted (IVW) method for exposures with more than one SNP [31]. Wald ratios (WRs) were derived for exposures with a single SNP [32]. Horizontal pleiotropy occurs when a SNP influences the outcome via a pathway other than the exposure of interest, this can bias estimation of the causal effect of an exposure and subsequently leads to type I statistical errors, thus violating a key assumption of MR (see **Supplementary Figure 1** [33]). MR methods which make differing assumptions regarding pleiotropy were performed as sensitivity analyses where genetic instruments were comprised of more than 3 SNPs: MR-Egger regression (each SNP is associated with the exposure independently of its pleiotropic effect), simple mode (across all SNPs effect estimates, the mode is 0), weighted mode (similar to simple mode, where weights are given to ratio estimates), and weighted median (at least 50% of all SNPs are valid instruments and no SNP contributes more than 50% of the weight; weight given to ratio estimates) [34–37].

While conventional MR methods assume effect homogeneity, large numbers of genetic instruments associated with an exposure can describe heterogenous effects, especially when there are multiple mechanisms through which the exposure might affect the outcome (e.g. variants associated with BMI may be associated with DVT via a number of alterations to the circulating proteome) [38]. To test for genetic heterogeneity, we used the maximum likelihood [39] estimator (fits a likelihood model to the summarized data, allowing for uncertainty in genetic associations with both the exposure and the outcome) and MR-Egger [35] for the exposures which were proxied by 2 or more variants.

#### Two-sample MR mediation analysis

The effect of BMI-associated proteins on DVT was estimated using the TwoSampleMR R package [30]. An IVW MR analysis was performed for FABP4, for which 3 SNPs were available to use as instruments. Wald ratios were derived for the remaining proteins. Where proteins were estimated to have a causal effect on DVT, a MR mediation analysis was performed to estimate the proportion mediated by a protein in the BMI-DVT link [40].

#### Multiple testing correction

As our MR-PheWAS estimated the causal relationship between a large number of exposures and DVT, we used PhenoSpD to estimate the number of independent traits in order to correct for multiple testing [41]. We used GWAS summary data describing the top 1000 associated SNPs for each exposure to create a phenotypic correlation matrix by Pearson correlation. This correlation matrix was used as an input for PhenoSpD to assess the number of independent exposures through matrix spectral decomposition [42,43]. As PhenoSpD is not able to assess the correlation between traits which come from different studies (e.g. BMI from GWAS “A” can’t be correlated with BMI from GWAS “B”), the number of independent variables resulting from the PhenoSpD analysis was likely overestimated. This will have elevated the risk of type 2 error because of the applied multiple testing correction based on the PhenoSpD analysis. Another cause of falsenegative findings arises from the limited power of some instruments. This discrepancy in power leads to a variation in significance of traits which are most likely correlated. For example, although we found many traits related to adiposity to be associated with DVT (e.g. BMI, weight, body fat percentage), exposures such as “Body fat” were not.

#### Beta coefficient transformation

Linear mixed model (LMM) methodology has gained popularity in GWAS due to its ability to control for population structure and deal with large datasets [44]. Regression coefficients are usually converted to odds ratios (ORs) or risk ratios (RRs) to make results interpretable. However, these cannot be calculated directly from LMM estimates and thus must be approximated. Using previously described methodology [45], we approximated logRRs for our LMM-derived MR estimates.

#### Bidirectional MR

Where there was evidence of an association with exposures tested in the MR-PheWAS, we performed a bidirectional MR analysis to assess the direction causality between a given exposure and DVT. This was conducted to identify potential pathways of reverse causation, which would invalidate MR assumptions [15].

### Conditional and colocalization analyses

Only one genetic instrument was available for some of the exposures investigated (N=10). As the Wald ratio estimator is susceptible to genetic confounding, we performed a conditional analysis for each single-SNP trait using the GCTA-COJO software [46] to identify any potential shared secondary signals in a 1MB region [47]. To perform these analyses, we downloaded summary statistics for these traits from OpenGWAS (https://gwas.mrcieu.ac.uk/) [48] and used genotypic data from the Avon Longitudinal Study of Parents and Children (ALSPAC) as a reference panel. Further details of the cohort are described elsewhere [49,50], in brief: 14,541 pregnant women with an expected delivery date of April 1, 1991, to December 31, 1992, were enrolled. We used the genotypic data of 8,890 mothers to perform our conditional analysis. Please note that the study website contains details of all the data that is available through a fully searchable data dictionary and variable search tool” and reference the following webpage (http://www.bristol.ac.uk/alspac/researchers/our-data/). Ethical approval for the study was obtained from the ALSPAC Ethics and Law Committee and the Local Research Ethics Committee.

Colocalization analysis uses Bayesian statistics to estimate whether an exposure and outcome share a causal signal in a region of the genome [51]. We used the R package “coloc” (https://cran.r-project.org/web/packages/coloc/) approximate Bayes factor (coloc.abf) function with default settings for prior probabilities to conduct a colocalization analysis with the following hypotheses: H0 (no causal variant), H1 (causal variant for trait 1 only), H2 (causal variant for trait 2 only), H3 (two distinct causal variants) and H4 (one common causal variant) [51]. We then used LocusZoom (https://locuszoom.org/) to provide visual evidence for the presence of a shared signal between our exposures and DVT.

## Results

### Associations of 973 exposures with DVT

With the aim to identify novel risk factors for DVT, we undertook a hypothesis-free MR-PheWAS to estimate the effect of 973 exposures with DVT. Of the 973 exposures investigated, 945 were identified as independent using PhenoSpD, setting the P-value threshold for our MR analysis at 5.43e-5. Fifty-seven exposures were estimated to influence DVT risk (**Figure 2**, **Table 1**). Sensitivity analyses results for all traits using additional MR methods are shown in **Supplementary Table 4**.

**Figure 2.**
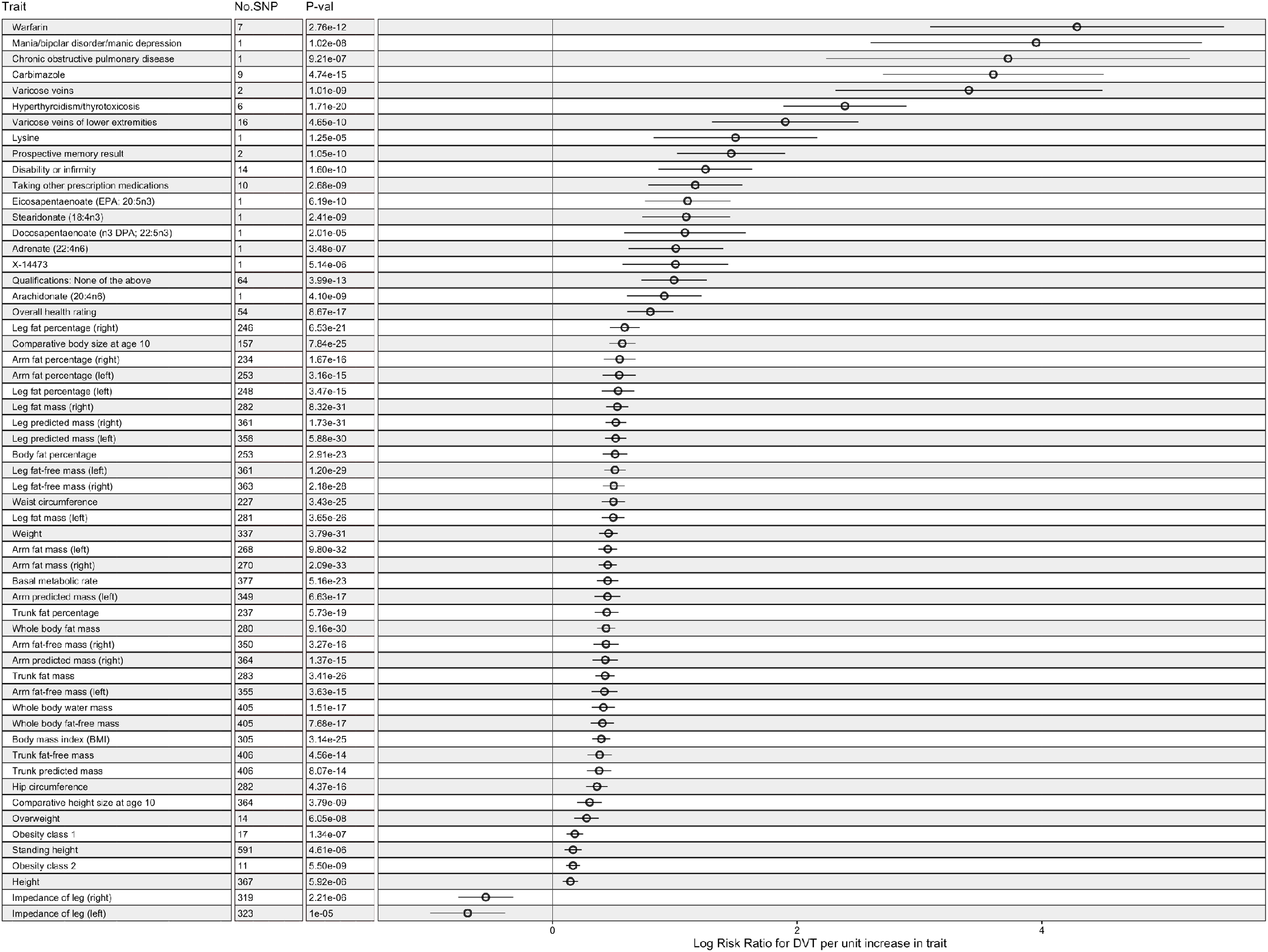
A many-to-one forest plot of the exposures which passed the P-value threshold following multiple testing correction (5.43e-5). Each trait is accompanied by two additional descriptive columns (No. SNPs and P-value), while log risk ratio (RR) is displayed to the right, alongside with the confidence intervals. MR methods: Inverse variance weighted (SNP > 1) and Wald ratio (SNP = 1).

We report several previously unidentified estimates, such as “Hyperthyroidism/thyrotoxicosis” (IVW Log RR: 2.39, 95% CI: 1.88 to 2.90; P = 8.69e-18), “Treatment/medication code: carbimazole” (IVW Log RR: 3.60, 95% CI: 2.70 to 4.50, P = 2.41e-12), “Chronic obstructive airways disease/chronic obstructive pulmonary disease (COPD)” (WR Log RR: 3.72, 95% CI: 1.39 to 4.37; P = 9.21e-07), “Varicose veins” (IVW Log RR: 1.90, 95% CI: 1.30 to 2.50; P = 2.36e-07), “Varicose veins of the lower extremities” (IVW Log RR: 3.40, 95% CI: 2.31 to 4.49; P = 5.13e-07) and “Mania/bipolar disorder/manic depression” (WR Log RR: 3.95, 95% CI: 2.60 to 5.30; P = 5.18e-06) (**Figure 2**, **Table 1**). We also report evidence for an effect of circulating fatty acids on DVT risk: “Eicosapentaenoate (EPA; 20:5n3)” (WR Log RR: 1.1, 95% CI: 0.75 to 1.45; P = 3.14e-07), “Stearidonate (18:4n3)” (WR Log RR: 1.09, 95% CI: 0.73 to 1.45; P = 1.22e-06), “Arachidonate (20:4n6)” (WR Log RR: 0.913, 95% CI: 0.61 to 1.22; P = 2.08e-06), “Adrenate” (WR Log RR: 1.01, 95% CI: 0.55 to 1.32; P = 3.48e-07), “Docosapentaenoate (n3 DPA; 22:5n3)” (WR Log RR: 1.08, 95% CI: 0.47 to 1.46; P = 2.01e-05) and the amino-acid “Lysine” (WR Log RR: 1.50, 95% CI: 0.61 to 1.96; P = 1.25e-05) with DVT (**Figure 2, Table 1**).

Adiposity, an established risk factor for DVT [52], and its related traits (N=24, see **Table 1** note) were all positively associated with DVT. These include traits found in previous MR studies, such as “Body Mass Index” (IVW Log RR: 0.40, 95% CI: 0.32 to 0.47; P = 1.60e-22), fat mass e.g. “Whole body fat mass” (IVW Log RR: 0.44, 95% CI: 0.36 to 0.51; P = 4.65e-27), fat-free mass e.g. “Whole body fat-free mass” (IVW Log RR: 0.41, 95% CI: 0.31 to 0.50; P = 3.90e-14) and fat percentage e.g. “Body fat percentage” (IVW Log RR: 0.51, 95% CI: 0.41 to 0.61; P = 1.48e-20) [53] (**Figure 2, Table 1**). A number of adiposity-related traits were found to be associated with DVT that have not been previously investigated in an MR framework, such as “Waist-hip ratio” (IVW Log RR: 0.50, 95% CI: 0.40 to 0.59; P = 1.74e-22), “Hip circumference” (IVW Log RR: 0.36, 95% CI: 0.28 to 0.45; P = 2.22e-13) and anatomically-specific measurements e.g. “Leg fat percentage (right)” (IVW Log RR: 0.59, 95% CI: 0.47 to 0.71; P = 3.32e-18) (**Figure 2, Table 1**). Another previously-associated trait is “Height” (IVW Log RR: 0.15, 95% CI: 0.08 to 0.21; P = 5.92e-06) [54]. Other associated height-related traits not previously investigated in an MR framework include “Standing height” (IVW Log RR: 0.17, 95% CI: 0.09 to 0.24; P = 4.61e-06) and “Comparative height size at age 10” (IVW Log RR: 0.30, 95% CI: 0.20 to 0.40; P = 1.93e-06) (**Figure 2, Table 1**).

Over 50% of the exposures (N=31) which passed our P-value threshold were found to have heterogenous effects between instruments using the maximum likelihood method. Of these, most (N=24) were traits related to body size (mass and adiposity). The remaining heterogenous traits were: basal metabolic rate (PHet: 3.71e-03); warfarin treatment (PHet: 5.66e-40); “Height” (PHet: 1.58e-03); “Standing height” (PHet = 4.61e-06); “Comparative height size at age 10” (PHet = 1.93e-06); “Impedance of leg (right)” (PHet: 4.23e-06) and “Impedance of leg (left)” (PHet: 9.96e-21). These findings were consistent with our IVW and MR-Egger heterogeneity analyses (**Table 1**).

MR-Egger estimates indicated strong evidence of horizontal pleiotropy for “Qualifications: None of the above” (intercept = −5.69e-04, P = 3.35e-02), “Impedance of leg (right)” (intercept = 2.58e-04, P = 3.22e-04) and “Impedance of leg (left)” (intercept = 2.22e-04, P = 7.24e-03) (**Table 1**). We were unable to assess whether the “Prospective memory result” trait was pleiotropic, as this exposure was instrumented using only 2 SNPs. In bidirectional MR analyses, DVT was estimated to increase warfarin treatment (“Treatment/medication code: warfarin” (beta = 0.29; SE = 0.02; P = 1.79e-30)), implying reverse causation (**Table 2**).

**Table 2.**
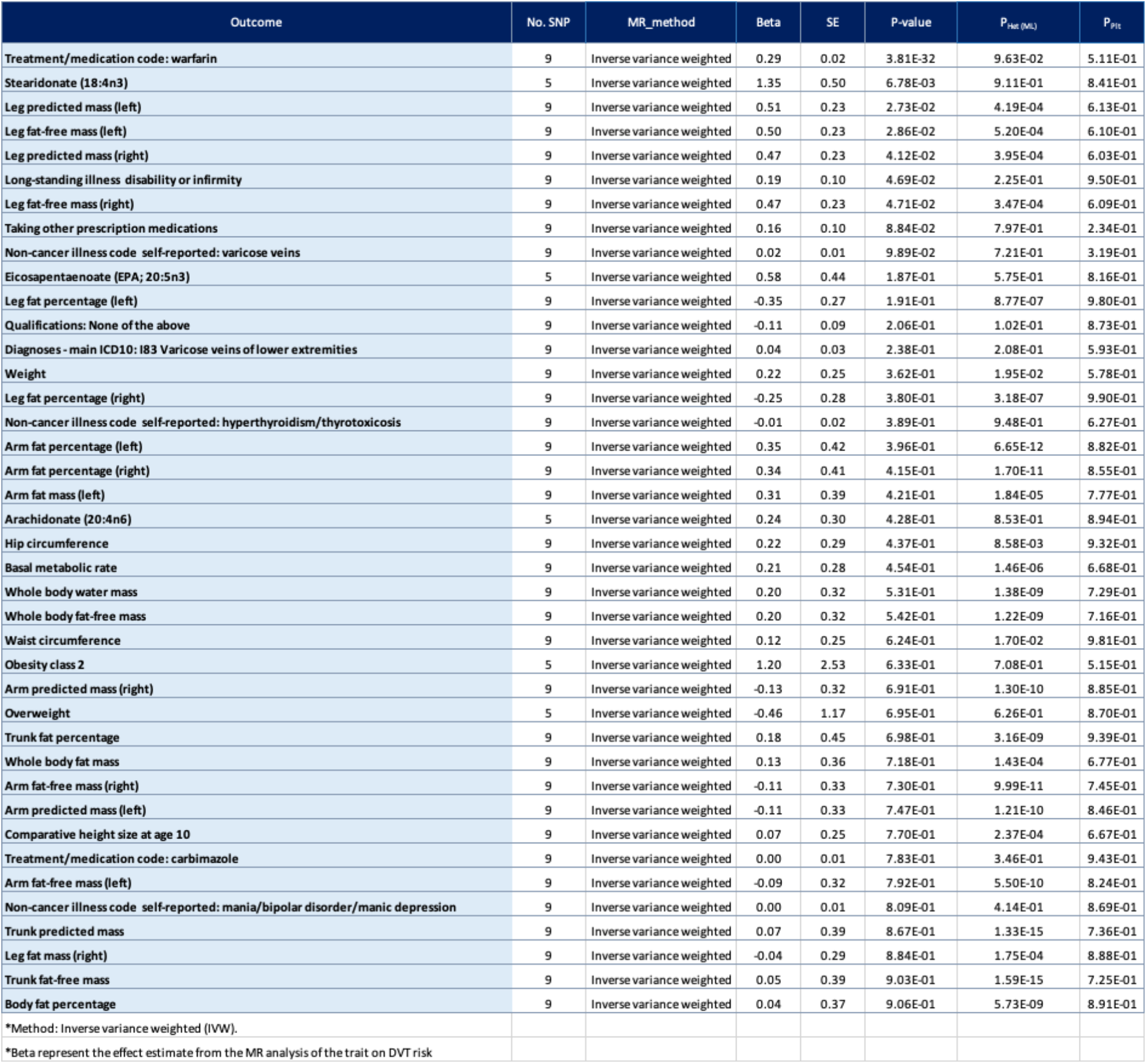
Reverse MR results of DVT on phenotypes from **Table 1**.

### Estimates of BMI-driven proteins with DVT

Of the 57 traits estimated to increase risk of DVT (**Table 1, Figure 2**), 24 were adiposity-related traits. While adiposity is an established risk factor for DVT, the biological mechanisms underlying the effect of adiposity and DVT are not well understood. To investigate the underlying mechanistic connection between adiposity and DVT we used a two-sample MR mediation analysis to test whether changes to levels of circulating blood proteins, driven by adiposity, were in part responsible for this association. Together, two recent MR studies have demonstrated that BMI causally affects the levels of 15 circulating proteins [18,19]. Our analyses provide evidence of a causal effect for 3 of these proteins on DVT risk: Neurogenic locus notch homolog protein 1 (NOTCH1; WR Log RR: 0.57, 95% CI: 0.45 to 0.68; P = 1.12e-23), Plasminogen activator inhibitor-1 (PAI-1; WR Log RR: 0.42, 95% CI: 0.30 to 0.54; P = 4.27e-12) and Inhibin beta c chain (INHBC; WR Log RR: −1.18, 95% CI: −2.18 to −0.69; P = 0.002). Mediation analysis was performed for PAI-1 (the only protein where BMI-protein and protein-DVT effect estimates were consistent in directionality, as otherwise the proportion of the effect mediated would be a negative number): the proportion of the BMI-DVT effect mediated by PAI-1 was estimated to be 18.56% (**Table 3, Figure 3, Supplementary Table 3**).

**Figure 3.**
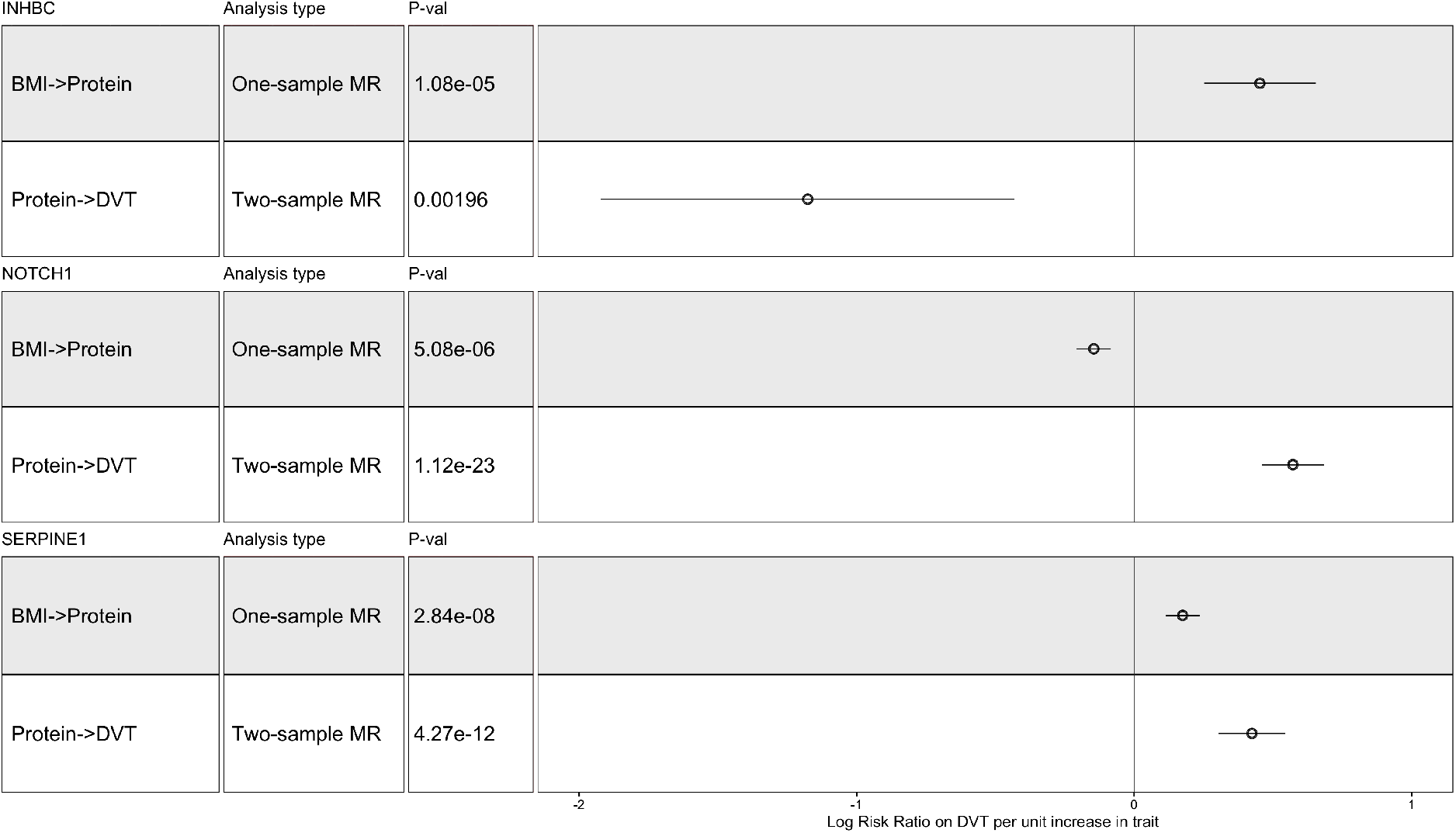
A many-to-one forest plot of the three BMI-associated proteins which passed the multipletesting corrected P-value threshold (0.003) in the MR analysis. Each protein is accompanied by two additional descriptive columns (type of analysis conducted and P-value), while the effect is displayed to the right, alongside with the confidence intervals (Beta coefficient/Log RR ± 95% CI). Effect sizes of BMI on proteins taken from Goudswaard et al [18] and Zaghlool et al [19].

**Table 3.** Mediation MR analysis results between BMI-associated proteins and DVT. The indirect effect was first calculated, followed by derivation of the proportion mediated.

| Table 3 |  | MR analysis of BMI-associated protein levels on DVT. Two-step MR of the indirect effect of BMI on DVT through protein levels.<br>Multiple-testing corrected P-value threshold: 0.0028 |  |  |  |  |  |  |  |  |  |  |  |
| --- | --- | --- | --- | --- | --- | --- | --- | --- | --- | --- | --- | --- | --- |
| Exposure | Protein abbreviation | Author | Year | PMID | MR_method | Log Risk Ratio* | CI (95%) |  | SE | P-value | No. SNP | Beta coefficient - BMI to protein estimate* | Proportion (%) mediated by protein |
| Neurogenic locus notch homolog protein 1 | NOTCH1 | Sun BB | 2018 | 29875488 | Wald ratio | 0.57 | 0.45 | 0.68 | 0.057 | 1.12E-23 | 1 | -0.15 | Effect not consistent |
| Plasminogen activator inhibitor 1 | PAI-1 | Sun BB | 2018 | 29875488 | Wald ratio | 0.42 | 0.30 | 0.54 | 0.061 | 4.27E-12 | 1 | 0.17 | 18.56 |
| Inhibin beta C chain | INHBC | Sun BB | 2018 | 29875488 | Wald ratio | -1.18 | -2.18 | -0.69 | 0.380 | 1.96E-03 | 1 | 0.45 | Effect not consistent |
| *LogRiskRatio is the logged value of the beta coefficient of the MR analysis into risk ratios. It can be read as an increase in the LogRisk of DVT per increase in circulating protein levels. |  |  |  |  |  |  |  |  |  |  |  |  |  |
| *BMI-Protein MR effect estimates from Goudswaard et al (https://doi.org/10.1038/s41366-021-00896-1) and Zaghlool et al (https://doi.org/10.1038/s41467-021-21542-4) |  |  |  |  |  |  |  |  |  |  |  |  |  |

Several of the proteins considered in our MR analyses could be instrumented using only one genetic variant, and therefore required a conditional and colocalization analysis to provide additional evidence of causality. There were no secondary signals after conditioning on the top SNP for each exposure-DVT pair. There was evidence of a shared causal variant for PAI-1 (PP.S = 97.5%), strengthening the evidence that there is a true causal relationship between the levels of this protein and DVT (**Table 4, Figure 4**).

**Figure 4.**
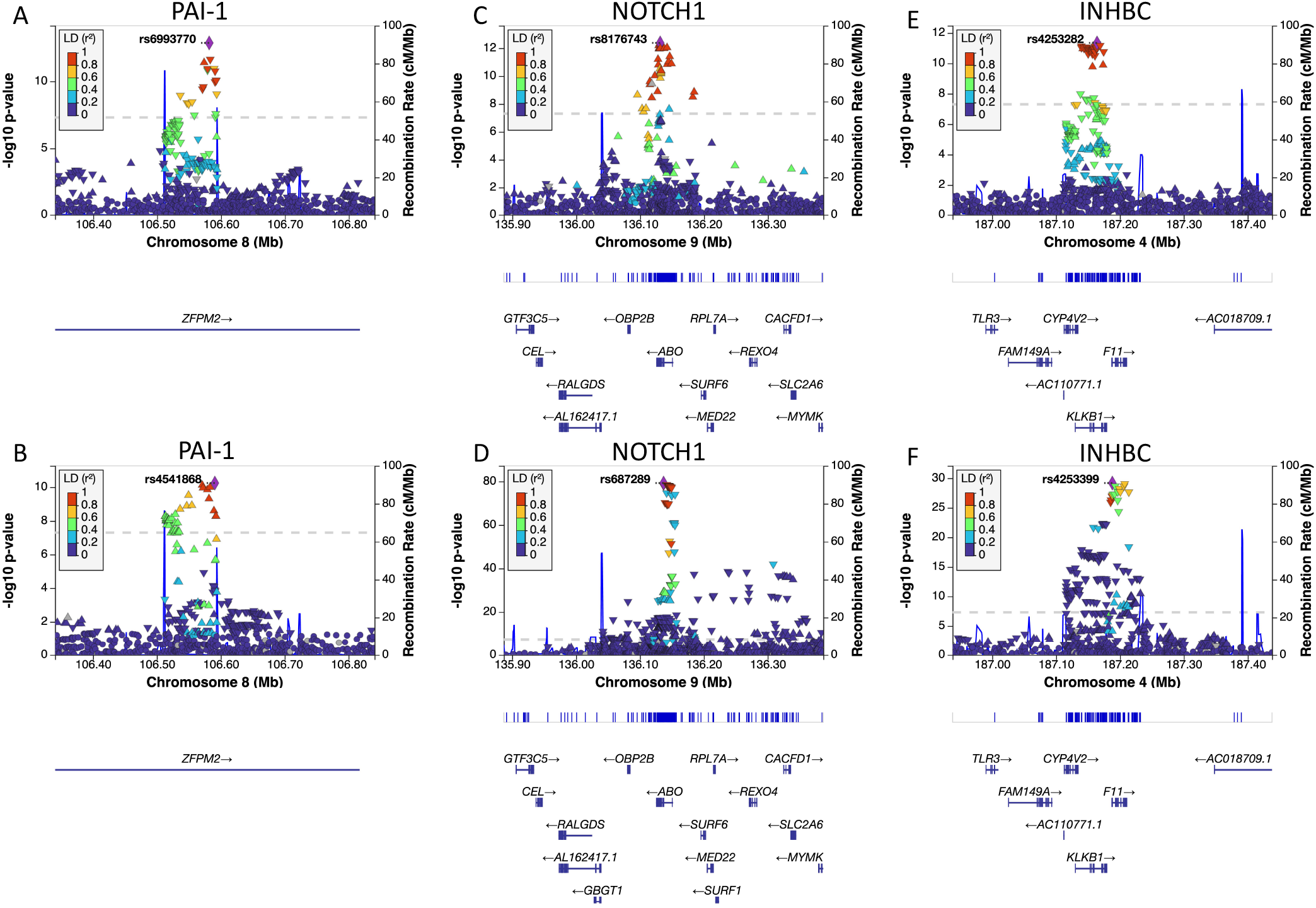
LocusZoom plots in a 1Mb region of the SNP used to proxy each protein in both exposure (A,C,E) and outcome (DVT: B,D,F) data: PAI-1 (A,B), NOTCH1 (C,D), INHBC (E,F). The top signal in the region is labelled in each figure. The x-axis represents the position within the chromosome, while the y-axis is the −log_10_ of the P-value. Each dot is a SNP, and the colours indicate how much LD there is between the reference SNP and the other genetic variants.

**Table 4.** Colocalization analysis results for exposures instrumented through only one SNP.

| Table 4 |  | Fully Bayesian colocalisation results of prioritized traits on DVT. |  |  |  |  |  |
| --- | --- | --- | --- | --- | --- | --- | --- |
| Trait 1 | nr SNP | PP.H0 | PP.H1 | PP.H1 | PP.H3 | PP.H4 | PP.S* |
| Neurogenic locus notch homolog protein 1 | 3856 | 1.0694E-79 | 4.778E-73 | 2.2382E-07 | 0.99999972 | 6.0801E-08 | 0.00% |
| Inhibin beta C chain | 4079 | 1.1109E-29 | 2.6137E-23 | 4.2502E-07 | 0.99999948 | 9.3591E-08 | 0.00% |
| Lysine | 547 | 2.4588E-11 | 0.98338278 | 3.2772E-13 | 0.01310352 | 0.0035137 | 0.35% |
| Bipolar disorder / mania | 3533 | 0.47264348 | 0.43702738 | 0.03965284 | 0.03665077 | 0.01402554 | 1.40% |
| Chronic obstructive pulmonary disorder | 4229 | 0.0766326 | 0.83975957 | 0.00333097 | 0.03645779 | 0.04381907 | 4.38% |
| X-14473 | 655 | 6.292E-07 | 0.83967623 | 6.959E-08 | 0.09280245 | 0.06752062 | 6.75% |
| Docosapentaenoate | 614 | 1.9077E-08 | 0.62830917 | 1.1181E-09 | 0.03649044 | 0.33520037 | 33.50% |
| Adrenate | 626 | 1.8886E-18 | 0.5838098 | 1.1167E-19 | 0.03413747 | 0.38205274 | 38.20% |
| Stearidonate | 674 | 5.34E-11 | 0.50441818 | 3.2335E-12 | 0.03007888 | 0.46550294 | 46.60% |
| Eicosapentanoate | 633 | 2.8064E-17 | 0.22721212 | 1.6606E-18 | 0.01268473 | 0.76010315 | 76.00% |
| Arachidonate | 626 | 4.9721E-77 | 0.17796851 | 2.9399E-78 | 0.00971061 | 0.81232088 | 81.20% |
| Plasminogen activator inhibitor 1 | 2604 | 3.0254E-13 | 1.9614E-06 | 3.9637E-09 | 0.02472248 | 0.97527556 | 97.50% |

## Discussion

With the aim to identify novel causal risk factors for DVT, we performed a hypothesis-free MR-PheWAS of 973 exposures to DVT, of which 57 passed a conservative P-value threshold for evidence of causality. We confirmed causality for several previously established risk factors for DVT (such as BMI and height) and have identified several novel putative causal risk factors (such as hyperthyroidism and COPD). Of the 57 exposures estimated to influence DVT risk, 24 were adiposity-related traits, therefore, we investigated whether the impact of adiposity on DVT is mediated by circulating proteins, known to be altered by BMI [18,19]. Here, we provide novel evidence that the circulating protein (PAI-1) has a causal role in DVT aetiology and is involved in mediating the BMI-DVT relationship.

Height has been previously associated with increased DVT risk [63] and our results align with this finding. With increased height, a greater volume of blood is required which can increase the stress on blood vessels, disrupting haemostasis [63]. Fat-free mass was also estimated to increase risk of DVT in our study. While counterintuitive, this effect could be mediated through height, as taller people usually have higher fat-free mass [53,54]. As expected, many body size related traits showed evidence of heterogeneity, likely due to the large number of SNPs used to instrument these traits and the many underlying biological pathways explaining variation in adiposity.

Interestingly, we found an effect of the genetic predicator for warfarin treatment on DVT risk. Warfarin is an anticoagulant used to treat DVT which reduces the production of vitamin K-dependent proteins involved in coagulation (FVIIa, FIXa, FXa, and thrombin) [3]. It has been reported that initial warfarin dosage may result in skin necrosis and a hypercoagulable state due to reductions in protein C and protein S levels, paradoxically increasing the risk of DVT [64]. At the same time, the underlying need for warfarin could be causing DVT (i.e. the SNP is increasing the vitamin K epoxide reductase complex 1 (VKORC1) activity, thereby increasing synthesis of clotting factors). Our bidirectional MR provided evidence of reverse causality indicating individuals prescribed warfarin may already be suffering from a form of VTE.

Venous blood stasis caused by immobility is also a known risk factor for DVT [3]. Here, we report evidence that long standing illness, disability, or infirmity increases DVT risk. A proposed mechanism is stasis of blood flow in the veins which can be either due to a particular neurological condition or due to the paralysis of the lower limbs [65]. Immobility may also arise due to hospitalisation and surgery, advanced age or a prolonged work-, air travel-, computer-related immobility, which have also been associated with an increased risk of DVT [66,67].

Our study also provides evidence for novel DVT risk factors. Hyperthyroidism has previously been proposed to contribute to DVT, as indicated by a recent systematic review and meta-analysis of cohort studies showing association with DVT (RR: 1.33, 95% CI: 1.28 to 1.39; I^2^ = 14%) [68]. In the present study, we for the first time provide novel evidence for a causal effect of hyperthyroidism/thyrotoxicosis on DVT risk (IVW RR: 10.91, 95% CI: 3.97 to 18.17; P = 3.14e-25). The underlying mechanism is not fully understood but may involve thyroid hormones (THs) promoting a hypercoagulable state and venous thrombi formation, by increasing plasma concentration of factor VIII, fibrinogen, plasminogen activator inhibitor 1 and vWF [69]. TH T4 may also directly enhance platelet function through integrin α_v_β_3_ [70]. In addition, THs enhance basal metabolic rate (BMR) and thermogenesis, both of which affect body weight. Indeed, we found that an increase in basal metabolic rate is associated with DVT. While a higher BMR should lead to lower BMI and thus lower DVT risk, it is likely that our results may be explained by the hyperthyroidism-associated mechanisms outlined above. A genetic proxy for requiring a prescription of carbimazole was also associated with increased risk of DVT. Carbimazole is a thionamide drug which has been used to treat thyrotoxicosis for over 60 years. It reduces the levels of circulating TH by binding to thyroid peroxidase, the enzyme required for TH production. As we found hyperthyroidism/thyrotoxicosis to be positively associated with DVT, we would expect carbimazole to protect against it. However, our MR estimate predicted a positive estimate. This novel finding is likely to be due to selection biases of the DVT GWAS, as those who are taking medication against hyperthyroidism are more likely to suffer from DVT [71]. It does, however, strengthen the link between hyperthyroidism and DVT risk.

Our MR estimates also support evidence of a causal association between varicose veins and increased risk of DVT. A common occurrence in varicose veins is the impaired action of leaflet valves, which prevent the blood from flowing backwards. This results in the inability of the blood to fully return to the heart, leading to the enlargement of the veins, and in time, potentially an increased risk of DVT due to stasis [72]. Varicose veins have been outlined as a possible risk factor in general practice patients in Germany[73], as well as in a Chinese retrospective study of over 100K people [72].

COPD was also associated with an increased risk of DVT. COPD is a severe chronic respiratory disease, having been studied extensively for its role in PE [74]. Indeed, both PE and DVT are more prevalent and underdiagnosed in people with COPD [75]. We were unable to perform a sensitivity analysis, as only one SNP was used to instrument this trait, and our colocalization analysis did not provide evidence that would support our MR estimates. With the observational evidence present, future studies with more genetic variants to instrument this trait would be worthwhile.

Finally, as adiposity is an established risk factor for DVT, the estimates we observe between adiposity-related traits and DVT most likely reflect true causal relationships. The estimate we report here for BMI (RR: 1.49, 95% CI: 1.38 to 1.60; P = 3.14e-25) is consistent with a previous MR study conducted in individuals of Danish descent (OR: 1.57, 95% CI: 1.08 to 1.97; P = 3e-03) [10]. In addition, our results are in agreement with the estimated effect of BMI on VTE in the FinnGen consortium (MR RR: 1.58, 95% CI: 1.28 to 1.95; P = 2.00e-05) [53]. Higher adiposity is associated with dysregulated metabolism, which is one factor that can promote a hypercoagulable state and impair venous return, increasing the chance of thrombi formation [55]. Given that 42% of the traits we found to be associated with DVT were adiposity-related and we previously found that adiposity is associated with changes to the circulating proteome [18,19], we hypothesised that adiposity-driven changes to the circulating proteome may promote DVT. Our analysis revealed estimates for levels of 3 BMI-driven circulating proteins (NOTCH1, PAI-1 and INHBC) with DVT. However, only the estimates for PAI-1 were directionally consistent for a potential mediator of the BMI-DVT relationship. Circulating levels of PAI-1 were positively associated with BMI and with DVT. These results are consistent with the known role for PAI-1 in inhibiting fibrinolysis (breakdown of a clot) [56]. In addition, PAI-1 expression has been previously found to be associated with DVT formation in mice [56] and in humans after total hip arthroplasty [57].

Although estimates were inconsistent with it being a potential mediator of the BMI-DVT relationship, we found that circulating INHBC levels were negatively associated with DVT, suggesting it may have a protective effect. Inhibins are part of the growth and differentiation superfamily of transforming growth factor beta (TGF-β) [58] and play a role in inhibiting the levels of follicle-stimulating hormone (FSH) produced by the pituitary gland [59]. Although we did not find evidence of causality between FSH and DVT, a recent study showed that FSH can enhance thrombin generation [60]. This discrepancy could be due to INHBC acting through a different pathway compared to FSH.

Finally, although again inconsistent with it being a potential mediator of the BMI-DVT relationship, we found that higher NOTCH1 expression was associated with an increased risk of DVT. NOTCH1 plays a role in responses to microenvironmental conditions, vascular development and is a shear stress and flow sensor in the vasculature [61]. However, a recent study conducted in mice suggested that NOTCH1 may be protective, as overexpression of miR-5189-3p in DVT led to increased expression of NOTCH1 and an improvement of venous thrombosis [62]. One explanation for this is that NOTCH is shed in the plasma and therefore expression levels within endothelial cells might be reduced.

In summary, we here confirmed estimates of previously identified traits on DVT (e.g. adiposity-related, height), and identified novel estimates (e.g. hyperthyroidism, COPD and varicose veins) with the disease. We also provide evidence that the relationship between adiposity and DVT is mediated by dysregulated levels of circulating proteins (PAI-1). These findings improve the understanding of DVT aetiology and have notable clinical significance, particularly in regard to hyperthyroidism and PAI-1.

## Availability of data and materials

GWAS summary statistics data are available on the MR-Base database. Scripts used to perform the analyses in this study are available on request.

## Contributions

AC, CJB, IH and EEV conceived the idea for the paper. AC conducted the analysis. All authors contributed to the interpretation of the findings. AC, LG, CJB, IH and EEV wrote the manuscript. All authors critically revised the paper for intellectual content and approved the final version of the manuscript.

## Acknowledgments

AC acknowledges funding from grant MR/N0137941/1 for the GW4 BIOMED MRC DTP, awarded to the Universities of Bath, Bristol, Cardiff and Exeter from the Medical Research Council (MRC)/UKRI. NJT is the PI of the Avon Longitudinal Study of Parents and Children (Medical Research Council & Wellcome Trust 217065/Z/19/Z) and is supported by the University of Bristol NIHR Biomedical Research Centre (BRC-1215-2001). NJT acknowledges funding from the Wellcome Trust (202802/Z/16/Z). EEV, CJB, and NJT acknowledge funding by the CRUK Integrative Cancer Epidemiology Programme (C18281/A29019). NJT, BE, EEV and CJB work in a unit funded by the UK Medical Research Council (MC_UU_00011/1 & MC_UU_00011/4) and the University of Bristol. EEV and CJB are supported by Diabetes UK (17/0005587) and the World Cancer Research Fund (WCRF UK), as part of the World Cancer Research Fund International grant program (IIG_2019_2009). LJG is supported by a BHF Accelerator Award Transition Fellowship (AA/18/1/34219). IH acknowledges funding by the BHF (PG/16/3/31833 and PG/16/21/32083). JZ is supported by Shanghai Thousand Talents Program and the National Health Commission of the PR China. BE acknowledges funding from Our Future Health, a company limited by guarantee registered in England and Wales (number 12212468) and a charity registered with the Charity Commission for England and Wales (charity number 1189681) and OSCR, Scottish Charity Regulator (charity number SC050917). We are extremely grateful to all the families who took part in this study, the midwives for their help in recruiting them, and the whole ALSPAC team, which includes interviewers, computer and laboratory technicians, clerical workers, research scientists, volunteers, managers, receptionists and nurses. The UK Medical Research Council and Wellcome (217065/Z/19/Z) and the University of Bristol provide core support for ALSPAC. This publication is the work of the authors and EEV and IH will serve as guarantors for the contents of this paper. This research was funded in whole, or in part, by the Wellcome Trust (217065/Z/19/Z & 202802/Z/16/Z). For the purpose of Open Access, the author has applied a CC BY public copyright licence to any Author Accepted Manuscript version arising from this submission. A comprehensive list of grants funding is available on the ALSPAC website (http://www.bristol.ac.uk/alspac/external/documents/grant-acknowledgements.pdf); This research was specifically funded by Wellcome Trust WT091310. This work was also supported by the Elizabeth Blackwell Institute for Health Research, University of Bristol, and the Wellcome Trust Institutional Strategic Support Fund (ISSF 204813/Z/16/Z). The funders of the study had no role in the study design, data collection, data analysis, data interpretation or writing of the report.

## Author information

### Affiliations

**MRC Integrative Epidemiology Unit at the University of Bristol, Bristol, UK**

Andrei-Emil Constantinescu, Caroline J. Bull, Nicholas J. Timpson, Emma E. Vincent

**Bristol Medical School, Population Health Sciences, University of Bristol, Bristol, UK**

Andrei-Emil Constantinescu, Caroline J. Bull, Nicholas J. Timpson, Emma E. Vincent

**School of Translational Health Sciences, University of Bristol, Bristol, UK**

Andrei-Emil Constantinescu, Caroline J. Bull & Emma E. Vincent

**School of Physiology, Pharmacology and Neuroscience, University of Bristol, Bristol, United Kingdom**

Lucy J Goudswaard, Samantha F Moore, Ingeborg Hers

**Department of Endocrine and Metabolic Diseases, Shanghai Institute of Endocrine and Metabolic Diseases, Ruijin Hospital, Shanghai Jiao Tong University School of Medicine, Shanghai, China.**

Jie Zheng

**Shanghai National Clinical Research Center for Metabolic Diseases, Key Laboratory for Endocrine and Metabolic Diseases of the National Health Commission of the PR China, Shanghai National Center for Translational Medicine, Ruijin Hospital, Shanghai Jiao Tong University School of Medicine, Shanghai, China.**

Jie Zheng

**Our Future Health. Registered office: 2 New Bailey, 6 Stanley Street, Manchester M3 5GS.**

Benjamin Elsworth

## Ethics declarations

### Ethics approval and consent to participate

UK Biobank received ethical approval from the NHS National Research Ethics Service North West (11/NW/0382; 16/NW/0274) and was conducted in accordance with the Declaration of Helsinki. All participants provided written informed consent before enrolment in the study. Ethical approval for ALSPAC was obtained from the ALSPAC Ethics and Law Committee and the Local Research Ethics Committees. Consent for biological samples has been collected in accordance with the Human Tissue Act (2004). Informed consent for the use of data collected via questionnaires and clinics was obtained from participants following the recommendations of the ALSPAC Ethics and Law Committee at the time.

### Consent for publication

All authors consented to the publication of this work.

## Supplementary information

### Additional file 1

**Supplementary Figures. S. Figure 1**, Mendelian randomization (MR) assumptions; **S. Figure 2**, Many-to-one Forest plot of the BMI-associated proteins which passed the P-value threshold after multiple testing correction; **S. Figures 3-5**, LocusZoom plot of the 1MB region within the top SNP for the proteins which did not pass the multiple testing adjusted P-value threshold.

### Additional file 2

**Supplementary Tables. S. Table 1**, Traits considered as exposures in the analysis of BMI-associated proteins on DVT; **S. Table 2**, Traits considered as exposures in the hypothesis-free MR analysis; **S. Table 3**, MR analysis of BMI-associated protein levels on DVT; **S. Table 4**, Secondary hypothesis-free analysis of traits on DVT with additional MR methods (where possible).

