## Supplementary Figures for "A phenome-wide approach to identify causal risk factors for deep vein thrombosis"

Pg. 1

**Supplementary figure 1. Mendelian randomization (MR) assumptions.** MR works in a similar way to a randomized controlled trial, exploiting the essentially random allocation of alleles at conception and the independent assortment of parental variants at meiosis. MR uses genetic variants (G) as proxies (instruments) to investigate whether an exposure (E), is causally associated with a disease outcome (O), in this case DVT. E is causally associated with O if the following conditions are held: (1) the genetic variant (G) is a valid instrument, in that it is reliably associated with E; (2) there is no independent association with O, except through E; and (3) the instrument is independent of any measured or unmeasured confounding factors (C).

Pg. 2

**Supplementary figure 2. Many-to-one Forest plot of the BMI-associated proteins which passed the P-value threshold after multiple testing correction.** Each protein is accompanied by four additional descriptive columns (GWAS author, MR method, No. SNPs and P-value), while log risk ratio (RR) is displayed to the right, alongside with the confidence intervals. MR methods: Inverse variance weighted (SNP > 1) and Wald ratio (SNP = 1).

Pg. 3-5

**Supplementary figures 3-5. LocusZoom plot of the 1MB region within the top SNP for the proteins which did not pass the multiple testing adjusted P-value threshold.** The top signal is displayed on the left for the pQTL data and on the right for the DVT data. The x-axis represents the position inside the chromosome, while the y-axis is the -log10 of the P-value. Each dot is a SNP, and the colours indicate the amount of LD between the reference SNP (top signal in the region) and the other genetic variants.

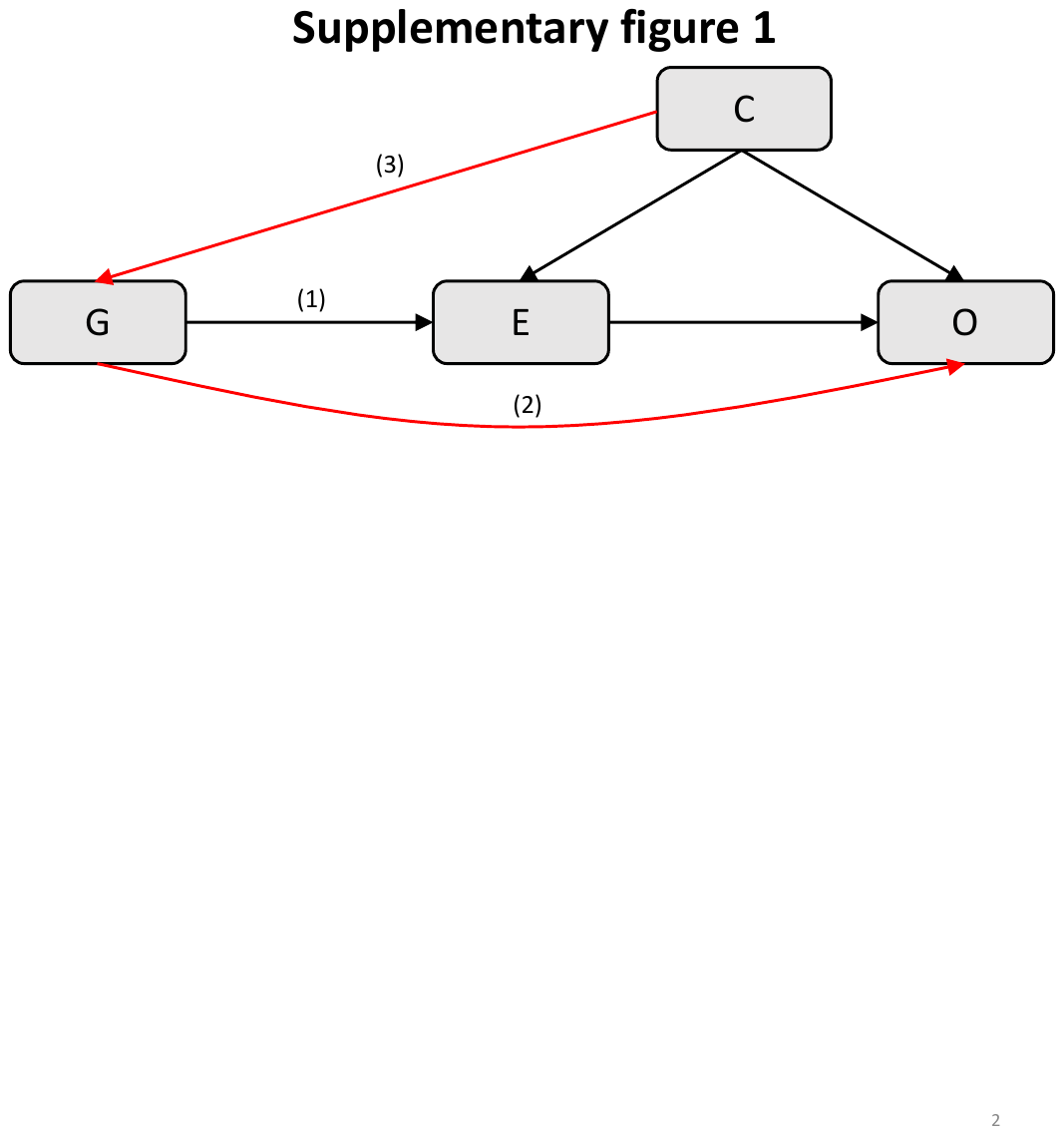

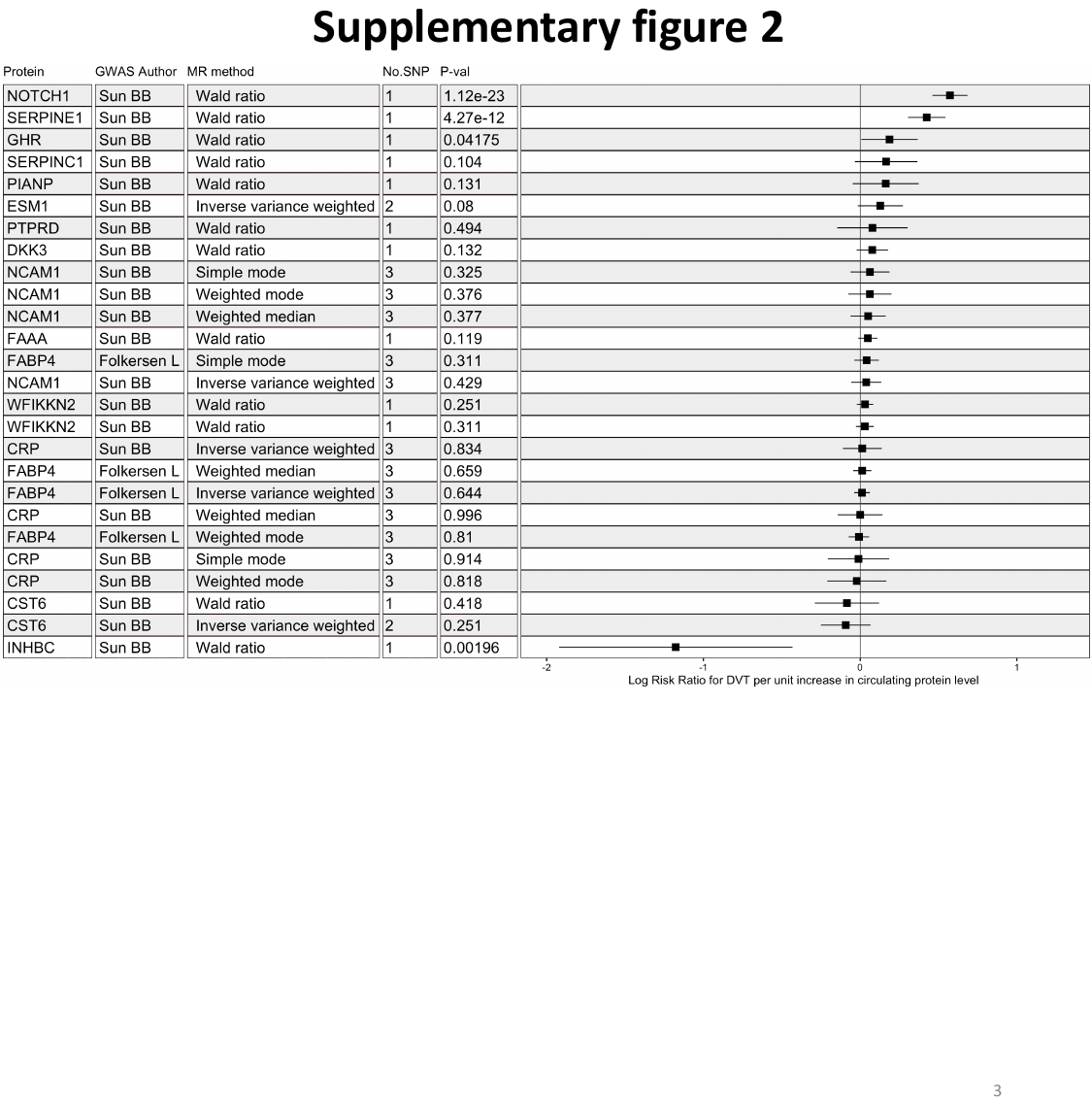

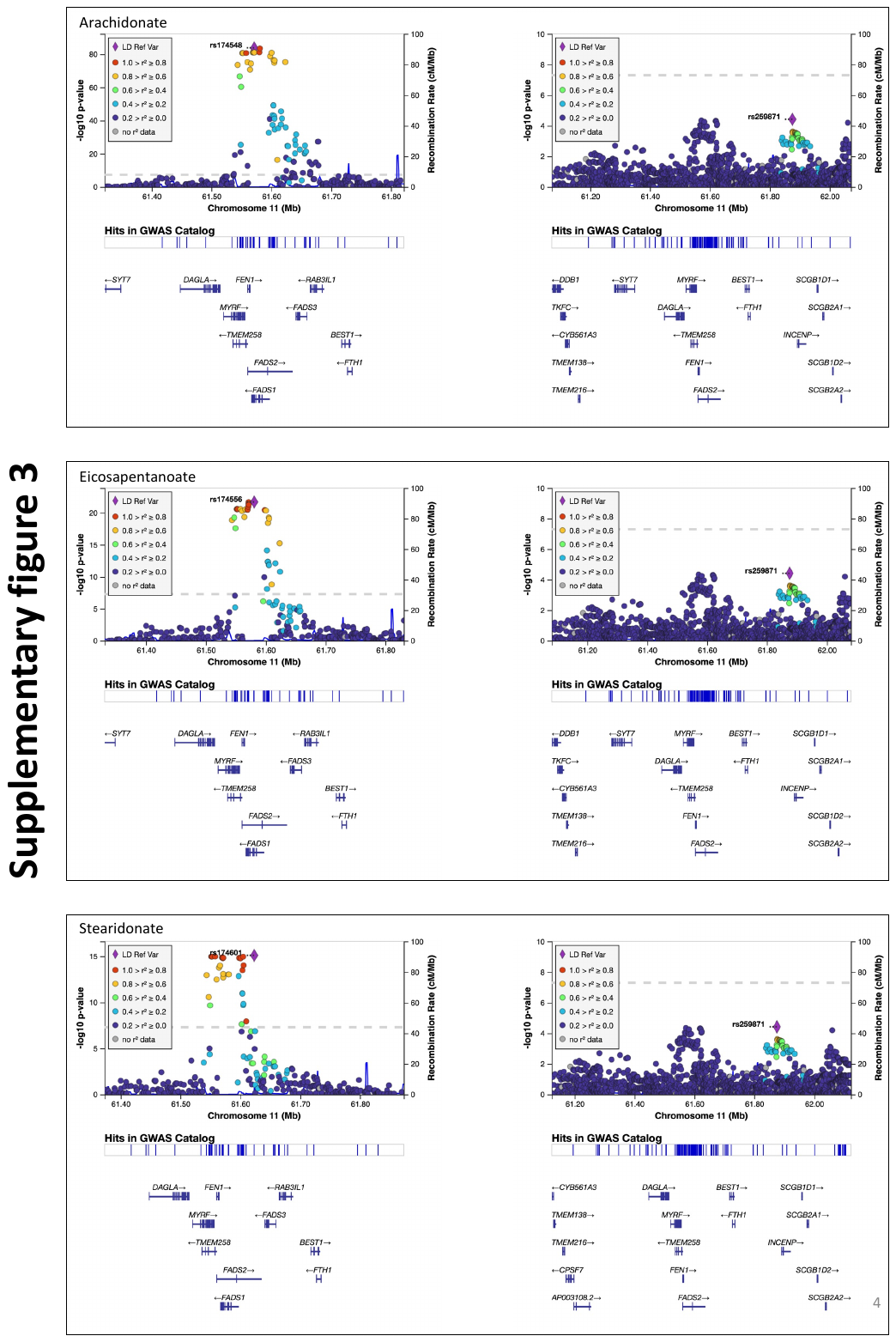

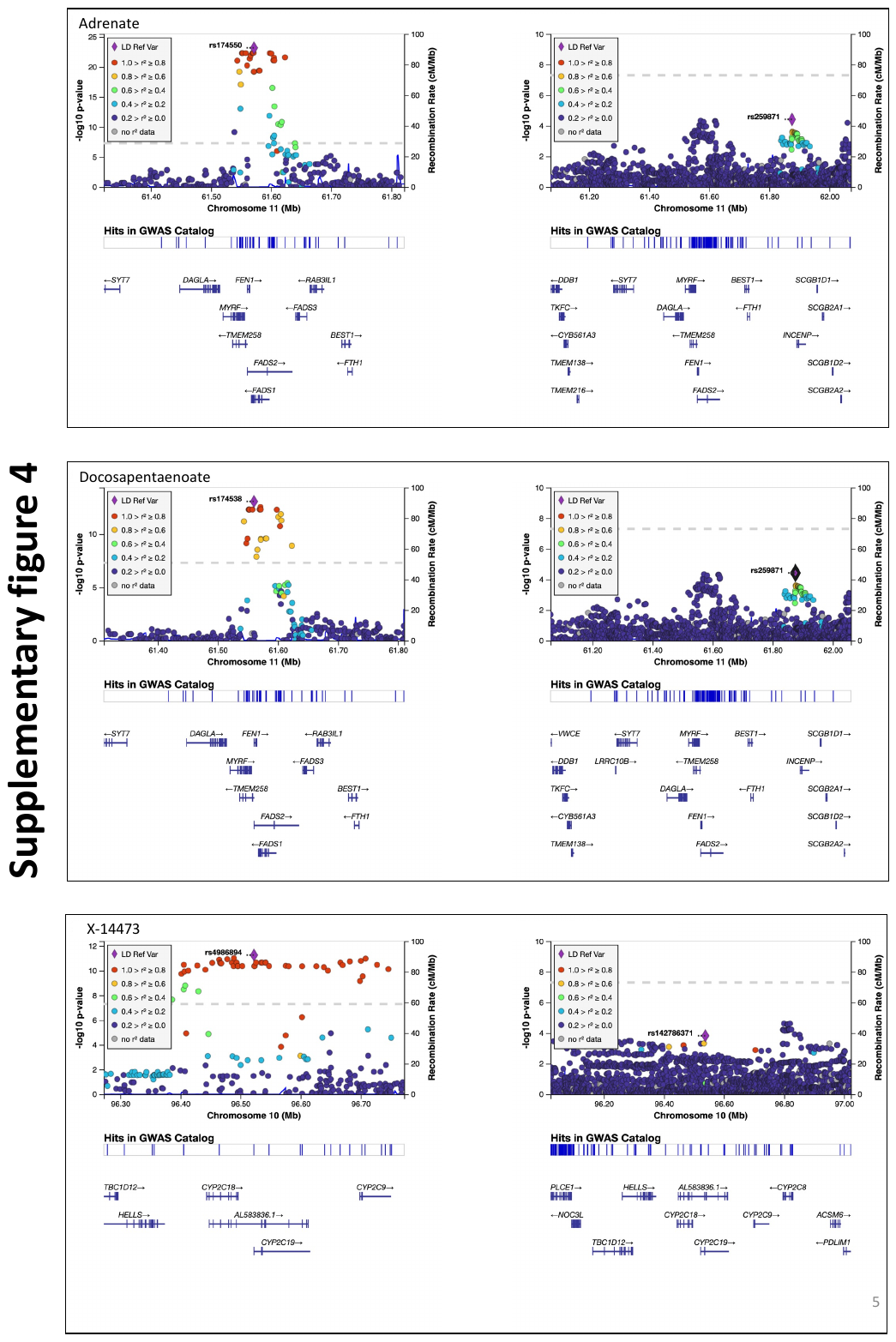

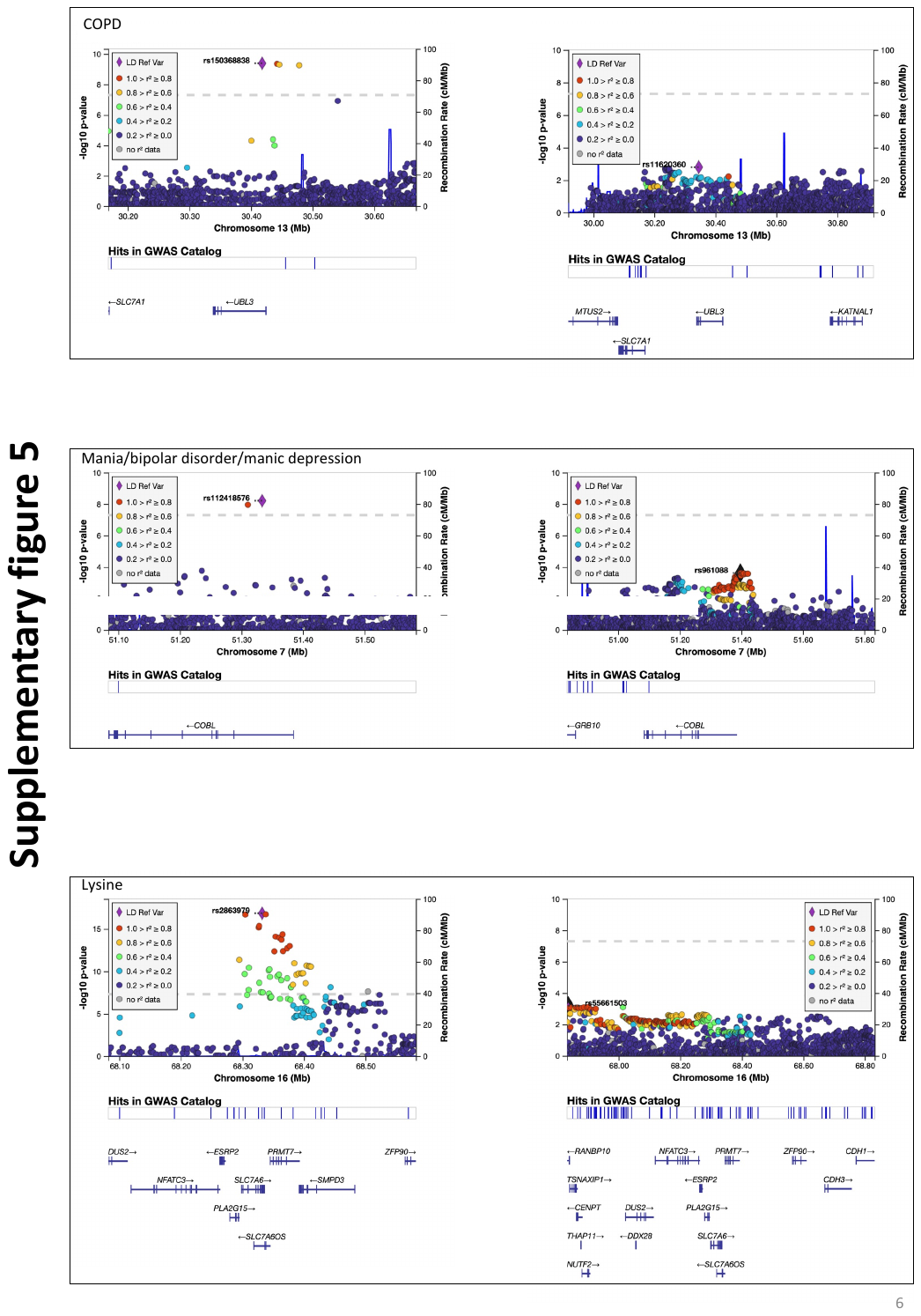
